# CTCF/cohesin organize the ground state of chromatin-nuclear speckle association

**DOI:** 10.1101/2023.07.22.550178

**Authors:** Ruofan Yu, Shelby Roseman, Allison P. Siegenfeld, Son C. Nguyen, Eric F. Joyce, Brian B. Liau, Ian D. Krantz, Katherine A. Alexander, Shelley L. Berger

**Author notes:** **One Sentence Summary:** CTCF and cohesin establish ground association between chromatin and nuclear speckles, impacting gene inducibility.

## Abstract

The interchromatin space in the cell nucleus contains various membrane-less nuclear bodies. Recent findings indicate that nuclear speckles, comprising a distinct nuclear body, exhibit interactions with certain chromatin regions in a ground state. Key questions are how this ground state of chromatin-nuclear speckle association is established and what are the gene regulatory roles of this layer of nuclear organization. We report here that chromatin structural factors CTCF and cohesin are required for full ground state association between DNA and nuclear speckles. Disruption of ground state DNA-speckle contacts via either CTCF depletion or cohesin depletion had minor effects on basal level expression of speckle-associated genes, however we show strong negative effects on stimulus-dependent induction of speckle-associated genes. We identified a putative speckle targeting motif (STM) within cohesin subunit RAD21 and demonstrated that the STM is required for chromatin-nuclear speckle association. In contrast to reduction of CTCF or RAD21, depletion of the cohesin releasing factor WAPL stabilized cohesin on chromatin and DNA-speckle contacts, resulting in enhanced inducibility of speckle-associated genes. In addition, we observed disruption of chromatin-nuclear speckle association in patient derived cells with Cornelia de Lange syndrome (CdLS), a congenital neurodevelopmental diagnosis involving defective cohesin pathways, thus revealing nuclear speckles as an avenue for therapeutic inquiry. In summary, our findings reveal a mechanism to establish the ground organizational state of chromatin-speckle association, to promote gene inducibility, and with relevance to human disease.

## Introduction

Nuclear bodies, defined as membrane-less, morphologically distinct sub-structures within the cell nucleus, have been implicated in chromosome organization (*1*) and gene regulation (*2*). Nuclear speckles are one type of nuclear body, comprised mainly of RNA and proteins involved in numerous aspects of RNA expression, including transcription, splicing, polyadenylation, RNA modification and export (*3*). In recent years, evidence has emerged to propose an active role for nuclear speckles in gene regulation. For example, some viruses utilize nuclear speckles to promote rapid production of mRNA (*4*), and, more generally, certain genes move closer to speckles in various biological processes such as heat shock (*5*), erythropoiesis (*6*), and p53 activation (*7*). Such increased chromatin-speckle association correlates with enhanced gene induction, potentially by bringing genes in close proximity to RNA biogenesis machinery (*8*).

Several studies indicate that the ground-state chromatin-nuclear speckle association exhibits a similar pattern across different cell types (*9*, *10*). In untreated cell lines, regions proximal to nuclear speckles (compared to nuclear speckle distal regions) are depleted of heterochromatic markers Lamin and H3K9me3, while enriched of gene-active markers H3K4me3 and H3K27ac; these regions also show higher gene density and overall higher gene expression levels (*9*, *11*). These findings suggest that association in the ground-state of nuclear speckles with chromatin may serve as a broad mechanism for gene regulation across various cell types. However, due to a lack of mechanistic knowledge of the factors that orchestrate the ground state chromatin-speckle association, the overall role of this layer of nuclear organization remains poorly understood.

Recent studies indicate that genes targeted to speckles upon induction are located in close proximity to nuclear speckles prior to induction. Pre-positioning at nuclear speckles was observed for genes activated and speckle-targeted by the transcription factor p53 (*7*), and for heat shock genes, which associate with speckles upon activation (*5*). These findings highlight that gene regulation by chromatin-speckle association is controlled on two levels: 1) inducible association upon change-of-state, and 2) basal association at the ground state, which then informs inducible association. Hence, there is evidence to support the existence of a key organizational principle of chromatin, that inducible genes are in close proximity to nuclear speckles. However, the mechanism that establishes this crucial basal state is unknown.

In this study, we investigated the mechanism governing the ground state of chromatin association with nuclear speckles. We identified the chromatin structural factors CTCF and cohesin as playing a vital role in establishing this association. Disruption of chromatin-speckle association led to impairment in the induction of speckle-proximal genes specifically. Moreover, we observed a similar disruption in chromatin-speckle association and impairment in gene induction in a rare genetic diagnosis, Cornelia de Lange syndrome, implying a broad relevance and function of ground state DNA-speckle association.

## Results

### CTCF depletion leads to dissociation between chromatin and nuclear speckle

To discover factors involved in association between nuclear speckles and chromatin at ground state, we utilized computational approaches to identify factors that displayed enrichment in genomic regions associated with speckles. Genome-wide datasets of DNA associated with speckles were previously generated using SON TSA-seq, a technique that uses proximity labeling to isolate DNA nearby the nuclear speckle protein, SON, which provides structure and scaffolding for the speckle (*7*, *9*, *10*). CTCF was reported to preferentially localize to chromatin associated with nuclear speckles [(*9*, *12*), and sample genomic regions shown in **Fig. 1A**]. However, this observed enrichment could be due simply to CTCF’s binding preference for active chromatin, which naturally tends to be in closer proximity to nuclear speckles compared to inactive chromatin. Indeed, we carried out correlation analysis of CTCF and found a genomic binding pattern that is comparable to the active chromatin marker H3K27ac, while showing an inverse relationship with the repressive marker H3K9me3 (**Fig.1A,1B**). Yet, upon comparing the ChIP-seq profiles of CTCF, its binding partner cohesin subunit RAD21 (*13*), and RNA Polymerase II (*14*), we discovered that CTCF and RAD21 genome binding exhibited greater similarity with SON than did PolII (**Fig.1A,1B**). This analysis suggests that enrichment of CTCF in speckle-associated DNA regions may surpass mere correlation with active chromatin.

**Figure 1.**
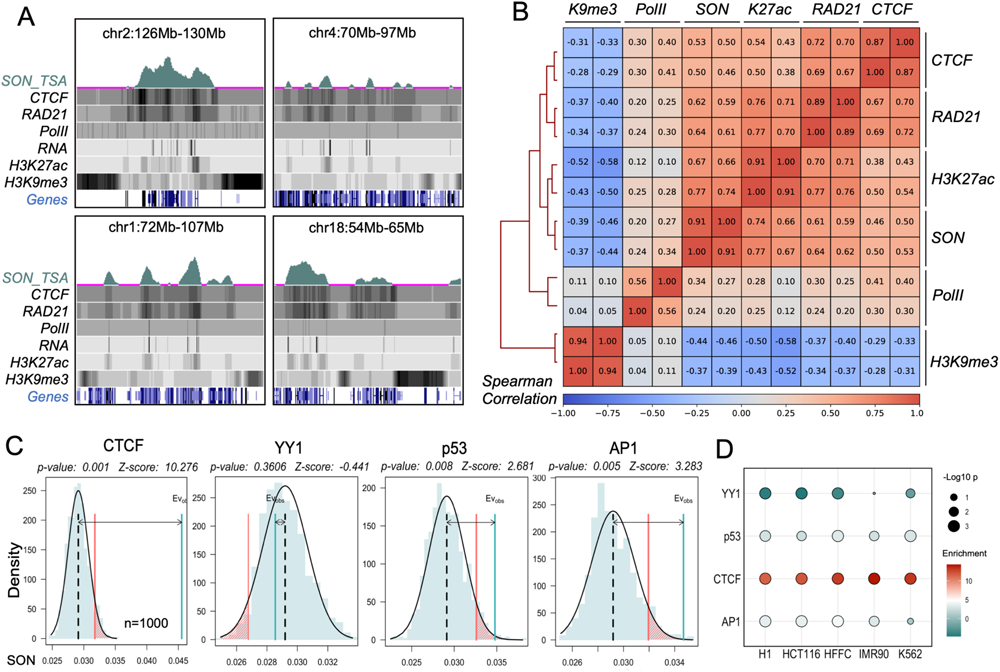
CTCF and cohesin colocalize with speckle associated regions genomically. A. Example signal tracks of SON-TSA seq and ChIP-seq of designated factors, histone modifications and RNA-seq, in IMR90 cells. B. Pairwise Spearman correlation between ChIP-seq signal of designated factors or histone modifications with SON-TSA seq, in IMR90 cells. C. The significance of enrichment of specific factors’ genomic binding sites within speckle-associated regions was assessed using permutation analysis. In the graph, the cyan shaded region under the curve represents the distribution of SON-TSA seq signal from 1000 randomly selected regions across the genome. The black dashed line indicates the median SON-TSA seq signal of all randomly selected regions. The red shaded fraction corresponds to the area with a significance level of p<0.05. The solid green line represents the median value of SON-TSA seq signal from all genomic binding sites of the designated factors. SON-TSA seq data from IMR90. D. Summary of permutation analysis results of designated factors in different cell-types.

To further test this hypothesis, we conducted a comparative analysis using a permutation test (*15*) to assess the degree of enrichment in speckle associated regions (SON TSA-seq) for multiple euchromatic factors, including another chromatin structural protein YY1 (*16*), and transcription factors p53 (*17*) and AP-1(*18*) in the fetal lung fibroblast cell line IMR90 (**Fig.1C**) and other cell lines with available SON-TSA seq data (*7*, *9*, *10*) (**Fig.1D**). We observed a notable and statistically significant enrichment of CTCF, supported by a high enrichment score (Z-score=10.276) and a minimal p-value (**Fig.1C, left**). This was in stark contrast to lower enrichment scores for YY1, p53 and AP-1 **(Fig.1C, right panels**). The results thus consistently demonstrated a pronounced and significant enrichment of CTCF binding sites within speckle-associated genomic regions across various cell lines. This enrichment exceeded the other factors, suggesting that CTCF’s presence in speckle-associated regions is unlikely solely to be attributable simply to its preferential positioning within active chromatin.

Given this correlative evidence, we next examined whether CTCF is necessary for association between chromatin and nuclear speckles. To profile the effect of reducing CTCF levels on DNA-speckle association genome-wide, we performed Cut&Run of the major speckle component, SON (*19*). We concluded that SON Cut&Run was a valid method for measuring the level of chromatin-speckle association based on previous publication (*20*) and the following quality-control results: (1) SON Cut&Run biological replicates exhibited good correlation (**Fig.S1A** and **S1B**); (2) SON Cut&Run signal exhibited a robust correlation with SON TSA-seq data, as observed in both the previously reported leukemia cell line K562 (*21*) and in our data obtained from IMR90 cells (**Fig.S1C**, SON TSA-seq data from (*7*); also see **Fig.2A**, compare TSA seq track to +siNeg); (3) Comparing SON Cut&Run to published large-scale multiplexed error-robust fluorescence *in situ* hybridization (MERFISH) data of speckles and DNA generated in IMR90 cells (*22*), we observed a clear inverse correlation between cytological DNA-speckle distance measured by MERFISH and SON Cut&Run signal (**Fig.S1D**), indicating that genomic locations with high SON Cut&Run signal are closer to nuclear speckle and vice versa; (4) Oligopaint DNA-FISH (*23*) of specific genes performed in our study also showed a similar inverse correlation as MERFISH (**Fig. S1E**). Taken together, these data establish SON Cut&Run as a reasonable technique for genome-wide measurement of DNA-nuclear speckle association.

**Figure 2.**
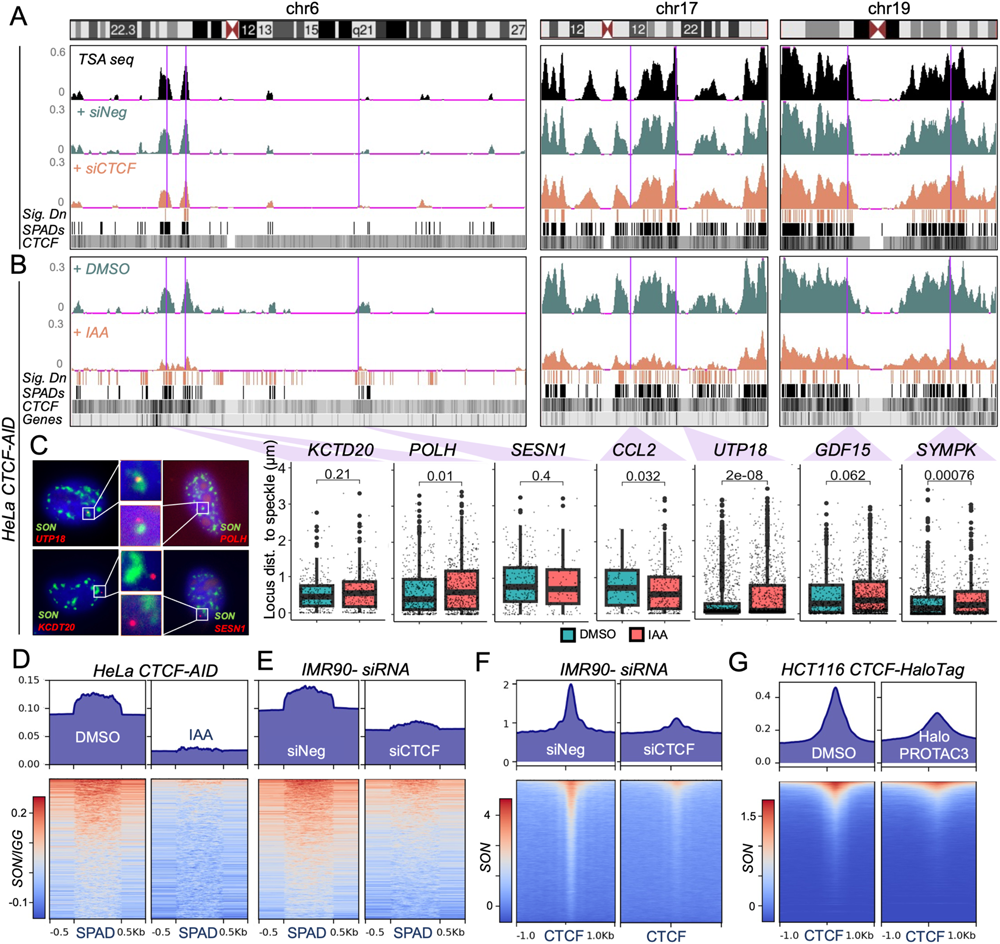
CTCF depletion leads to dissociation between chromatin and nuclear speckle. A. SON Cut&Run and CTCF ChIP-seq tracks under designated treatment at three representative chromosomes of IMR90 cells. SPADs track shows identified Speckle associated domains, Sig. Dn track shows regions with significantly decreased SON signal under CTCF depletion. B. SON Cut&Run and CTCF ChIP-seq tracks under designated treatment at three representative chromosomes of HeLa cells. C. (Left) Representative ImmunoDNA-FISH images with DAPI stained nuclei (blue), immunofluorescence of nuclear speckle component SON (green), and DNA probes targeting designated genes(red). (Right) Quantification of gene loci distance to nearest nuclear speckle upon DMSO (green) or auxin (red) treatment in CTCF-AID HeLa cell lines. All box plots show the median (center line), 1.5 times the interquartile range (whiskers), and the first and third quartiles (lower and upper hinges). Data points outside the first and third quartile range are displayed as outliers. D. Metagene plot and heatmap of IGG normalized SON Cut&Run signal across speckle associated domains (SPADs) in CTCF-AID HeLa cells with DMSO or 24hr auxin mediated knockdown of CTCF. E. Metagene plot and heatmap of IGG normalized SON Cut&Run signal at speckle associated domains (SPADs) in IMR90 cells, with 48hr siRNA knockdown of CTCF or negative control. F. Metagene plot and heatmap of SON Cut&Run signal across all CTCF binding sites in IMR90 cells, with 48hr siRNA knockdown of CTCF or negative control. G. Metagene plot and heatmap of SON Cut&Tag signal across call CTCF binding sites in HCT116 cells, with or without 32hr HaloPROTAC3 mediated knockdown of CTCF.

We employed two systems to profile genome-wide speckle association upon CTCF depletion: (1) primary lung fibroblast IMR90 cells with 48-hour *CTCF* siRNA knockdown and (2) a previously established auxin-inducible degron (AID) system in HeLa cells (*24*) with 24-hour auxin inducible degradation of CTCF (efficiency of CTCF knockdown validated in **Fig. S1F, G, H, I**). In both systems, an overall decrease of SON signal was detected in large regions that were speckle associated in the control condition (sample chromosomes shown in **Fig.2A** for IMR90 and **Fig.2B** for HeLa). To corroborate these findings for actual cytological distance, we performed Oligopaint DNA-FISH (*23*) to visualize eight genes located at diverse positions on four distinct chromosomes with varying nuclear speckle proximity measured by SON Cut&Run. The level of association between each gene (DNA-FISH probes designed across a 50kb region centered on the transcription start site) and nuclear speckles is represented by the distance between FISH foci of corresponding gene and the closest nuclear speckle, labelled by immunofluorescence against speckle marker protein SON (*19*) (examples shown in **Fig. 2C, left**). We observed a consistent pattern where genes with stronger association with speckles in the ground state (*UTP18*, *SYMPK*, *GDF15*, *POLH*) dissociate from nuclear speckles after a 24-hour auxin-induced degradation of CTCF (**Fig. 2C, right**). In contrast, genes with intermediate (*KCTD20*, *DDB2*) or low speckle proximity (*SESN1* and *CCL2*) showed lower dissociation from nuclear speckles (**Fig. 2C, right**; see *DDB2* in **Fig. S1J**). We noted as previously observed (*12*), depletion of CTCF did not overtly impact nuclear speckle morphology in size (**Fig. S1K**, Speckle Area), shape (**Fig. S1K**, Eccentricity and Extent), nuclear size (**Fig. S1K**, Nuclear Size), or average speckle count per nucleus (**Fig. S1K**, Speckle count). In conclusion, CTCF depletion led to a global dissociation between chromatin and nuclear speckle domains.

In further analysis, we examined changes in SON intensity within speckle-associated domains (SPADs, identified SPADs in control samples shown in **Fig.2A, 2B**), which are regions with significant SON enrichment. We identified a substantial decrease in SON levels across SPADs in both IMR90 and HeLa cell lines (**Fig. 2D**, **2E**). To better characterize the change in SON signal with regard to CTCF positioning, we analyzed SON signal level across all CTCF genomic binding sites within IMR90 cells and observed a drastic decrease upon siRNA mediated CTCF knockdown (**Fig. 2F**). To further explore CTCF’s role in speckle association across cell types, a third system was utilized. Here, HaloTAG-labeled CTCF underwent 32-hour degradation via HaloPROTAC3 in HCT116 colorectal carcinoma cells (*12*). We again observed an overt decrease of SON signal across CTCF binding sites upon CTCF depletion (**Fig. 2G**). Taken together, these findings support an important role of CTCF in mediating the ground-state of association between chromatin and nuclear speckles across multiple cell types.

### CTCF-cohesin interaction is necessary for chromatin-nuclear speckle association

One of the most intensively characterized function of CTCF is anchoring the cohesin complex. Over 90% of CTCF DNA-binding sites across the genome overlap with cohesin (*25*), and CTCF is known to stabilize cohesin on chromatin (*26*). CTCF binds to the genome through its zinc finger motif, and associates with RAD21 as well as the STAG1/ STAG2 proteins through its N-terminal region, facilitating the stabilization of cohesin at specific genomic sites (*13*, *25*). Given this known role of CTCF, we next investigated whether acute depletion of cohesin subunit RAD21 would dissociate chromatin from nuclear speckles. We utilized a RAD21-AID system in HeLa (*24*), where endogenous RAD21 is degraded rapidly (within 1 hour) after auxin treatment (validation in **Fig. S2A, B**). Similar to CTCF depletion (**Fig. S1K**), we found no overt change in nuclear speckle morphology with RAD21 degradation (**Fig. S2C**). Following acute RAD21 depletion for 1 hour, DNA-FISH indicated that genes located further away from speckles remained largely unaffected, while genes situated in close proximity to nuclear speckles at the ground state exhibited overall dissociation from nuclear speckles (**Fig. 3A**); these data closely matched the CTCF depletion (see **Fig. 2B**). In a genome-wide analysis using SON Cut&Run with IAA-induced RAD21 depletion, we found that SPADs showed global reduction of speckle association upon RAD21 loss (**Fig 3B**; see also tracks in **3A**, top), implicating the role of cohesin in mediating chromatin-speckle association. We also compared the SON intensity across all SPADs identified in either CTCF-AID or RAD21-AID controls and observed a similar decrease upon auxin treatment (**Fig. 3C**), suggesting that the depletion of CTCF and RAD21 leads to dissociation between speckles and chromatin at similar genomic regions.

**Figure 3.**
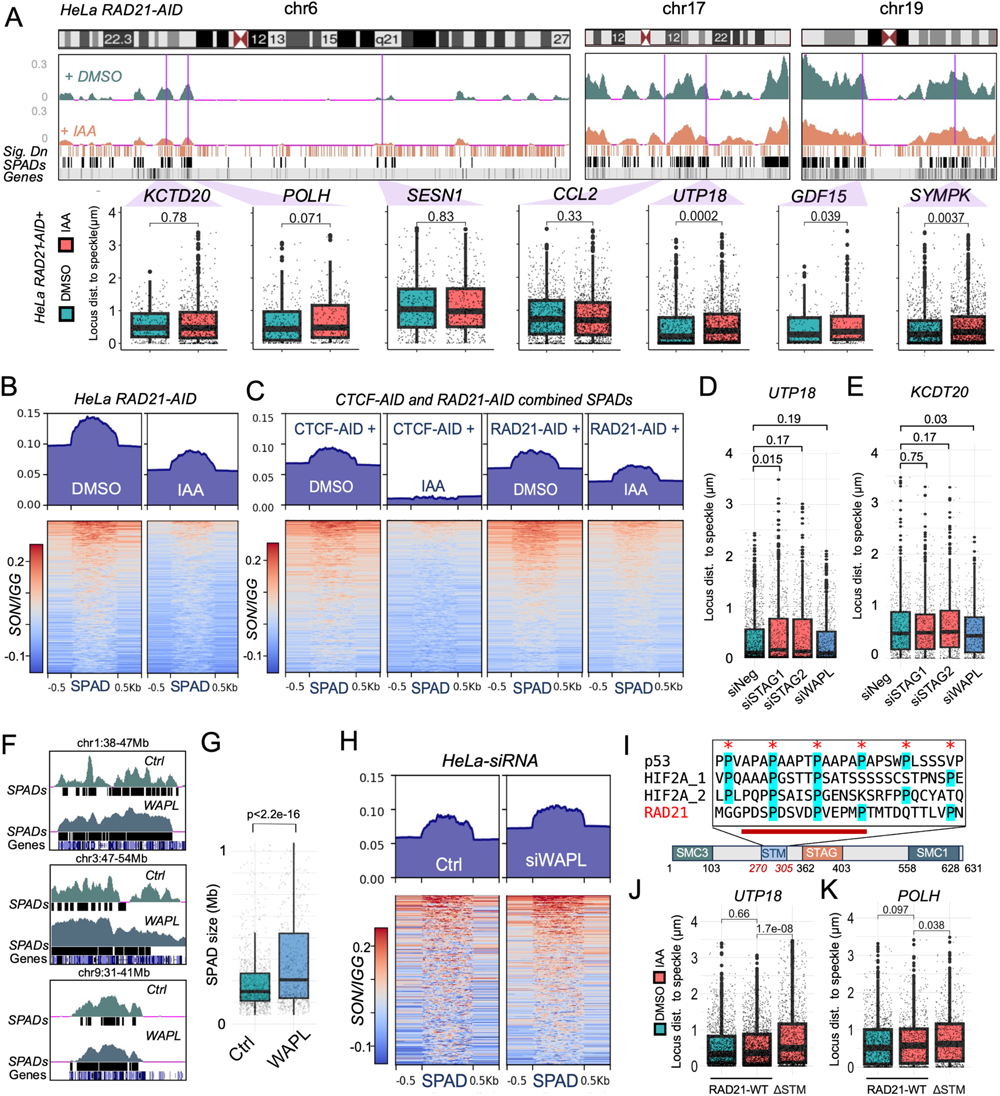
CTCF-cohesin interaction is essential for chromatin-nuclear speckle association. A. (Top) SON Cut&Run signal tracks under designated treatment at three representative chromosomes. (Bottom) Quantification of gene loci distance to nearest nuclear speckle upon 1hr DMSO (green) or auxin (red) treatment in RAD21-AID HeLa cells. B. Metagene plot and heatmap of IGG normalized SON Cut&Run signal at speckle associated domains (SPADs) in RAD21-AID HeLa cells with DMSO or 1hr auxin mediated knockdown of RAD21. C. Metagene plot and heatmap of IGG normalized SON Cut&Run signal at consolidated speckle associated domains (SPADs) of RAD21-AID and CTCF-AID cell line, under designated condition. D. Quantification of UTP18 gene loci distance to closest nuclear speckle upon siRNA knockdown of different cohesin pathway factors. E. Quantification of KCTD20 gene loci distance to closest nuclear speckle upon siRNA knockdown of different cohesin pathway factors. F. SON Cut&Run signal tracks at three representative regions, in HeLa cells with or without siRNA knockdown of WAPL. G. Comparison of SPAD sizes in Mb between HeLa cells with or without siRNA knockdown of WAPL. H. Metagene plot and heatmap of IGG normalized SON Cut&Run signal at speckle associated domains (SPADs), called in WAPL KD cells, in HeLa cells with or without siRNA knockdown of WAPL. I. (Top) Amino acids composition across putative Speckle Targeting Motif (STM) of designated factors. Asterisks represents periodic occurring prolines, red bar represents 15 amino acids targeted for deletion in ΔSTM mutant. (Bottom) Location of known RAD21 domains. J. Quantification of UTP18 gene loci distance to closest nuclear speckle in designated transfectants. K. Quantification of POLH gene loci distance to closest nuclear speckle in designated transfectants.

To further evaluate the role of the CTCF-cohesin interaction, we employed siRNA knockdown of cohesin subunits STAG1 and STAG2 (validation of knockdown in **Fig. S2D**), responsible for bridging CTCF-cohesin interaction (*13*). We found that *STAG1* or *STAG2* knockdown led to dissociation of the speckle proximal *UTP18* gene (**Fig. 3D**), whereas the gene *KCDT20*, positioned at an intermediate distance from the speckles, exhibited no notable alteration in response (**Fig. 3E**). Collectively, these results implicate the importance of the cohesin complex cooperating with CTCF in organizing chromatin around nuclear speckles.

Our findings strongly suggest that the loss of CTCF/cohesin results in decreased ground state chromatin-nuclear speckle association. To further test this hypothesis, we investigated whether, in contrast, gain of CTCF/cohesin leads to increased chromatin association with nuclear speckles. Cohesin bound to chromatin is unloaded by the releasing factor WAPL (*27*), hence, disruption of WAPL stabilizes cohesin binding on genome and antagonizes cohesin depletion (*28*). We utilized siRNA knockdown of *WAPL* in HeLa cells and performed DNA-FISH (validation of WAPL knockdown in **Fig. S2E**). We found no apparent change in gene-speckle distance of speckle-proximal gene *UTP18* (**Fig. 3D**), whereas the speckle-distal gene *KCTD20* showed increased speckle association in *WAPL* knockdown (**Fig. 3E**). From SON Cut&Run, we observed an apparent extension of SPADs following *WAPL* knockdown (sample regions shown in **Fig. 3F**, whole chromosome views in **Fig. S2F**, statistics shown in **Fig. 3G**), further corroborated by increase of SON signal across SPADs (**Fig. 3H**). Collectively, these findings indicate that the stabilization of cohesin through WAPL reduction expands genomic regions associated with speckles.

Given this possible role of cohesin in mediating chromatin-speckle association, we next investigated a potential domain within the cohesin subunit RAD21 required to target chromatin to speckles. A previous study from our lab identified a region within transcription factor p53 that is required for p53-mediated DNA-speckle association (*7*). This putative speckle targeting motif (STM) is characterized by periodicity of proline residues, which is also found in a second speckle-targeting transcription factor, HIF2A, and is highly enriched among gene regulatory proteins (*20*). We identified a similar region in RAD21 spanning M272 to N299 with proline periodicity (**Fig. 3I**). We assayed the function of this region to mediate chromatin-speckle association, by generating transient expression constructs of RAD21 with a 15aa deletion from Proline276 to Proline291 (**Fig. 3I, red bar.** Protein quantification in **Fig. S2G**). We note that this region and deletion does not overlap with known functional motifs within RAD21 (*29*). The wild type and deletion mutants were expressed in HeLa cells with auxin depletion of endogenous RAD21. We found that the wildtype RAD21 plasmid rescued speckle dissociation of *UTP18* (**Fig. 3J**) and *POLH* (**Fig. 3K**) genes upon auxin depletion, while the RAD21 ΔSTM mutant did not rescue speckle dissociation (**Fig. 3J** **and** **3K**). Based on these findings we propose the following model of CTCF/cohesin mediated chromatin-nuclear speckle association: CTCF anchors cohesin to chromatin, cohesin then brings chromatin and nuclear speckle to closer proximity via the STM region of RAD21.

### Disruption of nuclear speckle association hinders induction of nuclear speckle proximal genes

Previous studies have suggested that highly expressed genes often tend to be located in close proximity to nuclear speckles (*9*, *10*, *30*). We found, using gene ontology (GO) analysis of genes located in close proximity to speckles in both IMR90 and HeLa cells (genes with top 10% highest SON Cut&Run signal), a significant enrichment of “essential biological processes” (**Fig. 4A**). These processes included “gene expression” and various pathways of “macromolecule biosynthesis” (**Fig. 4A**). Hence, CTCF and cohesin through their role in mediating association between nuclear speckle and chromatin, may be of vital importance specifically in regulating certain crucial basal molecular pathways.

**Figure 4.**
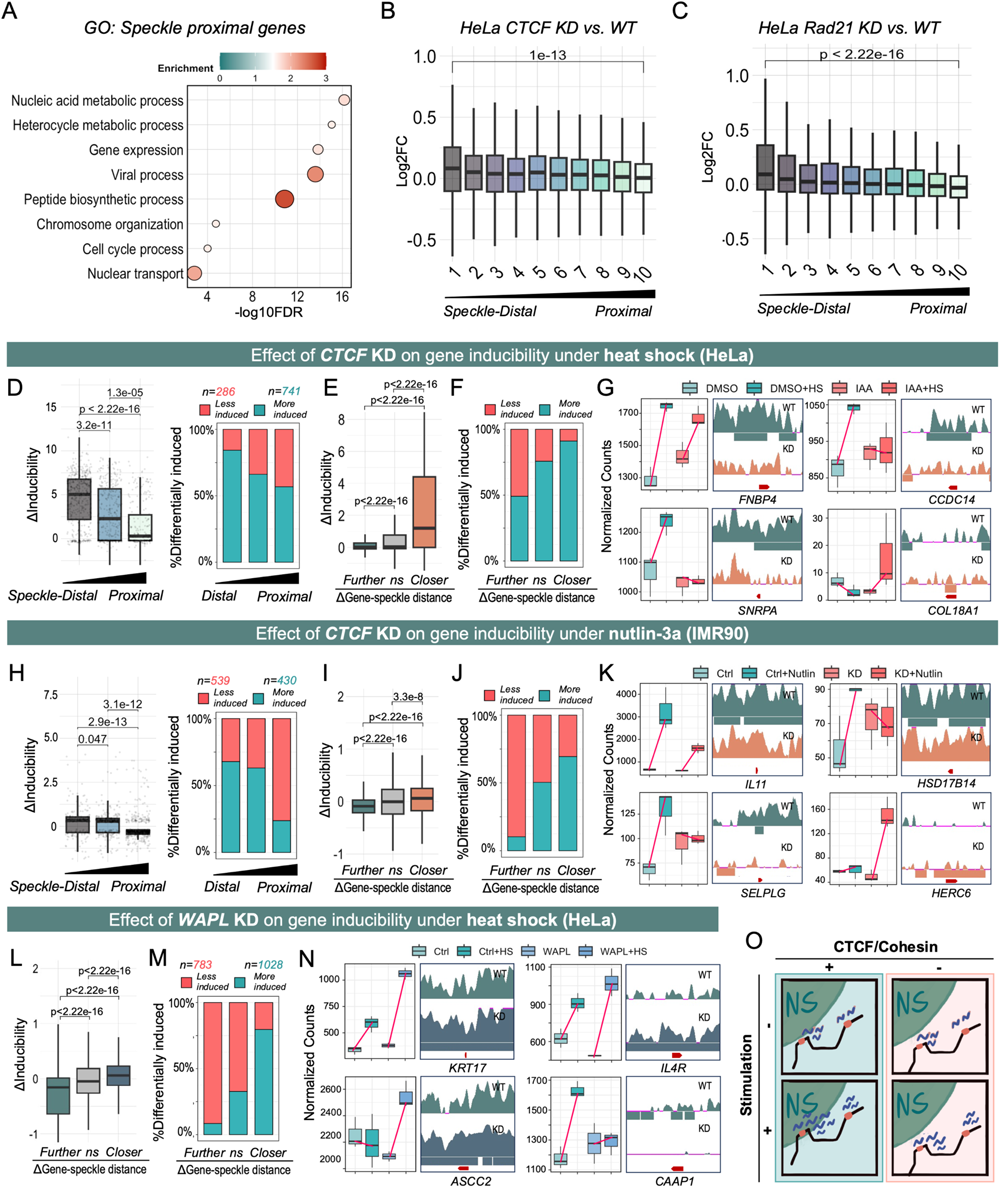
Disruption of nuclear speckle association hinders the induction of nuclear speckle proximal genes. A. Gene ontology (GO) categories enriched in nuclear speckle proximal genes in both IMR90 and HeLa cells. B. Log2 fold change between HeLa CTCF-AID cells with or without auxin mediated CTCF depletion for 24hr. Genes are binned according to their speckle proximity, calculated from SON Cut&Run signal over CDS. C. Log2 fold change between HeLa RAD21-AID cells with or without auxin mediated RAD21 depletion for 1hr. Genes are binned according to their speckle proximity, calculated from SON Cut&Run signal over CDS. D. Differential conditional effect (ΔInducibility) between auxin induced CTCF KD and DMSO treated HeLa cells under heat shock. Genes with significant conditional effect differences (p<0.05) were categorized into three bins based on nuclear speckle proximity. Left: Differential conditional effect represented by DeSeq2 calculated Log2 fold change. Right: Percentage of genes showing significantly higher (green) or lower conditional effect under heat shock. E. Differential conditional effects of genes with significantly different SON binding in auxin induced CTCF KD vs. DMSO treated control: Genes with reduced SON signal are labeled ’further,’ those with increased signal are labeled ’closer,’ and genes with insignificant signal changes are labeled ’ns’. F. Percentage of genes showing significantly higher (green) or lower conditional effect under heat shock, grouped by change in speckle proximity. G. Representative genes with significantly differential SON binding under auxin induced CTCF KD. Box plots represent RNA-seq normalized counts. In the SON Cut&Run track, the red arrow indicates the gene CDS location, and the regions underneath each track represent differential regions with preferential SON binding under the designated condition. H. Differential conditional effect between siRNA mediated CTCF KD and control IMR90 cells under nutlin-3a treatment. I. Differential conditional effects of genes with significantly different SON binding in siRNA mediated CTCF KD and control IMR90 cells. J. Percentage of genes showing significantly higher (green) or lower conditional effect under nutlin treatment, grouped by change in speckle proximity. K. Representative genes with significantly differential SON binding under siRNA mediated CTCF KD. L. Differential conditional effects of genes with significantly different SON binding in siRNA mediated WAPL KD and control HeLa cells. M. Percentage of genes showing significantly higher (green) or lower conditional effect under heat shock, grouped by change in speckle proximity. N. Representative genes with significantly differential SON binding under siRNA mediated WAPL KD in HeLa cells. O. Schematic model illustrating how changes in nuclear speckle proximity, resulting from CTCF or cohesin disruption, lead to differential conditional effects for speckle proximal and speckle distal genes.

Previous studies show that CTCF (*31*) and cohesin (*32*, *33*) perturbation impair activation of a subset of genes, and, importantly, both factors are essential as disruption of either can lead to developmental failure or cell death (*34–36*). In contrast, disruption of either CTCF or cohesin do not cause a broad change in steady state gene expression (*37–39*). Similarly, our RNA-seq of cells altering CTCF, RAD21, or WAPL levels showed only moderate changes of steady state gene expression (**Fig. S3A-C**). However, we grouped the transcriptome based on gene distance to speckles, and observed a small but significant downregulation of genes proximal to speckles in HeLa cells with auxin induced degradation of CTCF or RAD21 (**Fig. 4B, C**). These results indicate a potential role of CTCF/RAD21 in facilitating the expression of the gene subset located proximal to nuclear speckles, which our analysis above show is enriched in various vital biological processes (**Fig. 4A**).

Nuclear speckles are proposed to facilitate gene induction under various external stimuli, such as heat shock and nutlin-3a (*5*, *7*). Similarly, CTCF loss-of-function is more consequential for gene expression upon change of state (*31*, *39*). Thus, given the modest consequences of CTCF/cohesin perturbation on speckle associated gene expression at steady state (**Fig. 4B, 4C**), we next investigated gene activation under stress to determine whether CTCF/cohesin specifically facilitates induction of speckle-proximal genes. First we used heat as a stimulus, as heat shock is well-known to induce numerous protective genes including protein chaperones and DNA damage repair genes (*40*). We subjected HeLa cells to a heat shock of 1 hour at 45°C and detected induction of heat shock response genes in RNA-seq (Fold change >0 and p<=0.01, volcano plot in **Fig. S3D**). Using DESeq2 (*41*), we performed interaction analysis to examine the heat shock-specific effect of CTCF reduction on genes with diverse speckle proximity. All genes with significantly differential induction (p<0.05) were assigned to three bins, distal, medial, and proximal, based on SON Cut&Run signal. We found that CTCF depletion resulted in a drastic decrease in heat shock-dependent induction of speckle proximal genes compared to speckle distal genes, both in overall level (**Fig. 4D** **left**) and number of differentially induced genes within each bin (**Fig. 4D** **right**).

In a reciprocal analysis, we identified heat responsiveness of genes with significantly differential SON Cut&Run signal upon *CTCF* loss (as in (*7*)). Notably, genes that moved away from nuclear speckles (exhibiting decreased SON Cut&Run signal) showed lower heat shock induction both in terms of level (**Fig. 4E**) and number of genes (**Fig. 4F**) compared to genes that moved towards nuclear speckles. For example, *FNBP4*, *CCDC14*, and *SNRPA* genes each showed reduction of speckle association upon *CTCF* AID degradation and displayed lower gene induction by heat shock (**Fig. 4G**; compare difference between two blue bars and two pink bars; red line), while *COL18A1* showed a reverse trend (**Fig. 4G**). These findings indicate that, in response to heat shock, CTCF disruption has a heightened impact on the induction of genes located near nuclear speckles. Furthermore, increased proximity to nuclear speckles positively influences overall gene induction in response to heat shock.

Our data suggest that, by setting up ground-state chromatin-speckle association, CTCF/cohesin primes genes for induction. However, the effect of CTCF depletion on gene induction could also be due to its well-known role in DNA looping (for example, mediating enhancer-promoter loops (*39*)). To differentiate between these two scenarios, we utilized published CTCF ChIP and HiChIP data in HeLa cells (*42*). We found that, CTCF depletion hampers heat shock-induced activation of speckle proximal genes specifically (**Fig. S3E left**), while looping status showed minimal effect (**Fig. S3E right**). Therefore, gene induction defects due to CTCF disruption are more correlated with speckle proximity than is loop status.

A second instance of gene regulation by nuclear speckle association is gene activation by the p53 stress-responsive transcription factor (*7*). We assayed gene induction level in IMR90 cells with nutlin-3a treatment for 6 hours to activate the p53 signaling pathway, and compared reduction of CTCF. We found that, similar to heat shock in HeLa cells, siRNA-mediated *CTCF* knockdown led to a significant reduction of speckle-proximal gene induction compared to speckle-distal genes (**Fig. 4H**). Genes with decreased nuclear speckle association displayed reduced induction in both expression level (**Fig. 4I**) and quantity (**Fig. 4J**). For instance, *IL11*, *HSD18B14*, and *SELPLG* genes displayed reduced speckle association upon CTCF knockdown (**Fig. 4K**). They also exhibited lower gene induction under nutlin-3a treatment. Conversely, the *HERC6* gene, which demonstrated increased speckle association, showed heightened induction levels (**Fig. 4K**), underscoring the impact of CTCF disruption on speckle-proximal gene induction across multiple cell types.

To further corroborate the role of nuclear speckle proximity in gene induction, we compared RNA-seq results of heat-shocked HeLa cells with siRNA knockdown of *WAPL*, predicting that *WAPL* knockdown will increase gene expression of speckle associating genes. Consistently, we found that genes relocating towards nuclear speckles exhibited higher overall induction (**Fig. 4L**). Additionally, the majority of genes with significantly higher induction moved closer to speckles (**Fig. 4M**, example genes shown in **Fig. 4N**). Taken together, these findings provide numerous lines of evidence that, upon stimulation of a gene pathway, CTCF/cohesin-mediated speckle association acts to amplify induction of genes located in close proximity to speckles (**Fig. 4O**).

### Chromatin-speckle dissociation presents in CdLS patient derived cells with mutation of RAD21

Given the proposed roles of CTCF and cohesin in mediating association between chromatin and nuclear speckles, an intriguing question arises as to whether disease-associated mutations could alter DNA-speckle association. Variations in cohesin components RAD21 (*43*) and SMCs (*44*), and in the cohesin loading factor NIPBL (*45*) are causative for Cornelia de Lange Syndrome (CdLS), an autosomal dominant developmental disorder occurring in one per ten thousand births. CdLS is characterized in humans by microcephaly, small size, characteristic craniofacial dysmorphology, intellectual disability, gastrointestinal disorders, congenital heart defects, limb, and many other systemic differences (*46*). Although cohesin is vital for sister chromatid cohesion and cell cycle progression, surprisingly, structural chromatin defects are not overt in CdLS patient-derived cell lines(*47*, *48*). Since altered gene expression occurs in various models of cohesin mutations, developmental deficits in CdLS are proposed to arise from changes in gene expression, however, the mechanism is not fully understood (reviewed in *47*).

Given our findings that cohesin maintains DNA-speckle association and regulates gene inducibility, we investigated whether CdLS-linked mutations in RAD21 disrupt DNA-speckle association correlating with reduced gene induction. We utilized a CdLS patient derived Lymphoblastoid cell line (LCLs) bearing a duplication within the 5^th^ exon of RAD21: c.592_593dup; p. (Ser198Argfs*6); this mutation causes a premature stop codon that likely leads to nonsense mediated decay of the mutated transcript (*49*). We confirmed this genomic mutation by Sanger sequencing (**Fig. S4A**), and with RNA-seq showing ∼50% reduction of RAD21 transcript (**Fig. S4B**). Similar to cells bearing CTCF and RAD21 knockdown shown above (**Fig. S1K** and **Fig. S2C**), nuclear speckle morphology did not change overtly in patient LCL (P561) compared to LCLs derived from two healthy individuals (**Fig. S4C, S4D**). SON Cut & Run on patient derived LCLs revealed a striking reduction of genome-wide SON signal (**Fig. 5A, B**), indicating global dissociation between chromatin and nuclear speckles similar to CTCF or RAD21 knockdown in HeLa or IMR90 cells.

**Figure 5.**
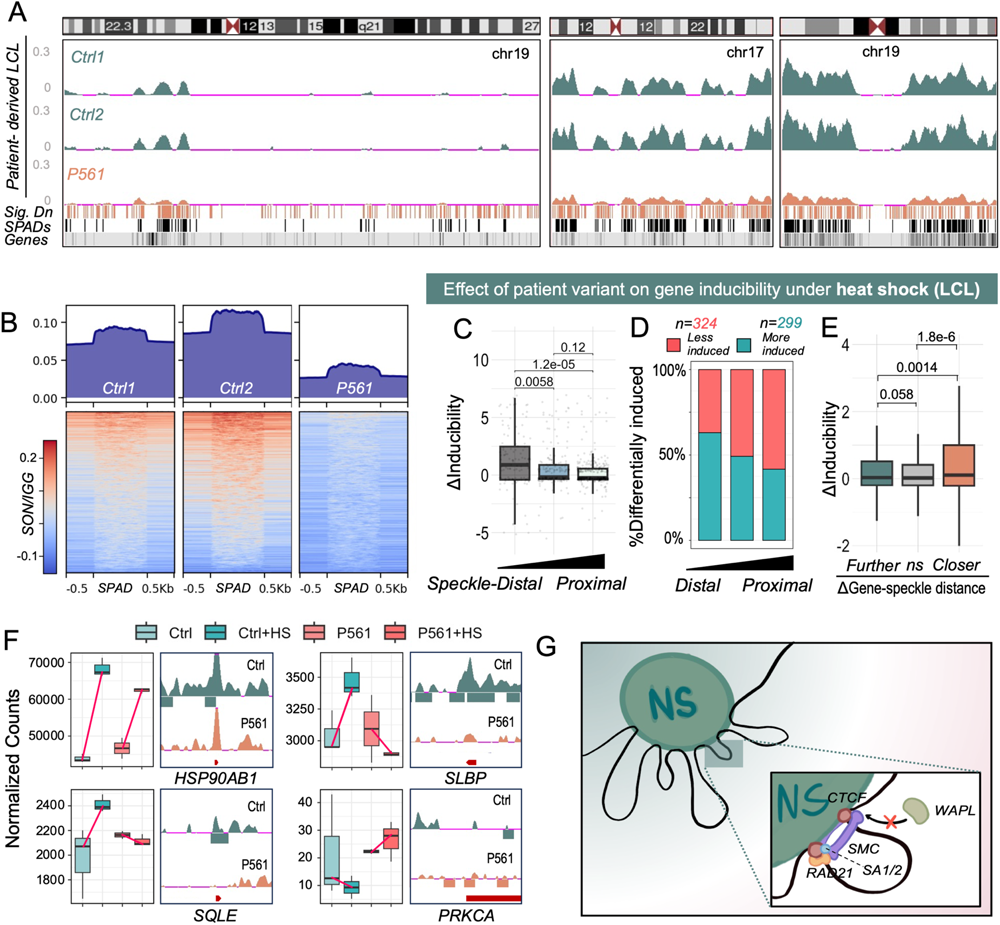
Chromatin-speckle dissociation presents in CdLS patient derived cell lines with mutation of RAD21 gene. A. Representative images with DAPI stained nuclei (blue), immunofluorescence of nuclear speckle component SON (green), in LCL cells from healthy donors (Ctrl1, Ctrl2) and patient 561 (P561-1, P561-2). B. SON Cut&Run tracks from designated healthy individual or patient derived LCLs at three representative chromosomes. C. Metagene plot and heatmap of SON Cut&Run signal across speckle associated domains (SPADs) in LCLs from designated healthy individual or patient. D. Differential conditional effects (ΔInducibility) between P561 and control LCL under heat shock. E. Differential conditional effects of genes with significantly different SON binding in P561 vs. control LCL. F. Representative genes with significantly differential SON binding in P561 cells. G. Schematic model: CTCF and cohesin associate genomic regions to nuclear speckle, enabling transcription factors (TFs) to bring nearby genes closer to nuclear speckle for expressional amplification upon activation.

To examine the effect of RAD21 mutation on gene induction, we heat-stressed patient-derived LCL cells for 1 hour and performed RNA-seq. Similar to CTCF disruption in HeLa and IMR90 cells (**Fig. 4**), we observed a notable decrease in the induction of genes located near speckles in P561 compared to healthy control (**Fig. 5C, 5D**), whereas the differential gene induction status between two control cell lines did not exhibit a clear correlation with nuclear speckle proximity (**Fig. S4E**). Additionally, genes with reduced speckle association in P561 were significantly more lowly induced compared to genes with increased speckle association (**Fig. 5E**). Among the genes exhibiting reduced induction (**Fig. 5F**), notable examples are the molecular chaperone gene HSP90AB1 (*50*) and stem loop binding protein SLBP (*51*) which is necessary for pre-mRNA processing of histone mRNA. Collectively, the disruption of CTCF/cohesin consistently affects the ground association between chromatin and speckles along with impaired induction of speckle proximal genes, both in laboratory cell lines and also, prominently, in patient cells derived from CdLS.

## Discussion

In this study, we present evidence supporting a new function of chromatin structural factors CTCF and cohesin in mediating the ground state of chromatin-speckle association. We found that disruption of CTCF or cohesin led to genome-wide dissociation between chromatin and nuclear speckles (**Fig. 2**, **Fig. 3**), and significantly impaired inducibility of genes located proximal to speckles (**Fig. 4**). Importantly, this effect was consistent across various cell lines. We propose that ground-state chromatin-speckle association primes genes for further conditional chromatin-speckle association under various stimuli, thereby amplifying expression of nuclear speckle-proximal genes upon induction (**Fig. 5G**). Moreover, we observed a similar disruption of chromatin-speckle association and impairment of gene induction in a cell line derived from a patient with Cornelia de Lange Syndrome, which notably involves RAD21 deficiency (**Fig. 5**), indicating that dysfunction of ground-state chromatin-speckle association occurs in human disease.

Our results provide a new perspective on gene regulatory roles of CTCF and cohesin. This is of particular importance considering that disruption of these factors does not induce drastic changes in the transcriptome under steady-state conditions (*37–39*). Rather, we find that depletion of CTCF and cohesin leads to impaired gene induction under ‘change-of-state’ conditions for a crucial gene subset, such as in response to heat-shock and p53 activation (**Fig. 4**), and our results point to this transcriptional effect as an outcome of CTCF/cohesin’s critical function in establishing the ground-state of chromatin-nuclear speckle association. Hence, we propose that the ground-state speckle association mediated by CTCF/RAD21 functions as a regulatory mechanism to boost gene expression in response to stimuli.

Both CTCF and RAD21 have established roles in facilitating chromatin loop formation. Consequently, further investigation is required to explore the relationship between nuclear speckles and chromatin conformation. It has been reported that nuclear speckle-associated chromatin shows enrichment in the A1 active subcompartment, and disruption of speckle protein SRRM2 reduces chromatin interactions in A1 (*52*). On the other hand, a recent study has reported minimal impact on chromatin loop formation upon disruption of nuclear speckles (*12*). These findings underscore the need for continued research to unravel the intricate interplay between nuclear speckles and the broader landscape of 3D chromatin organization.

Nuclear speckles are proposed to play a crucial role in facilitating gene expression (*5*, *7*, *8*). Despite the abundance of evidence supporting the correlation between speckle proximity and gene expression levels, a significant limitation in the field has been the absence of causative evidence due to the essential nature of speckle components, such as SON and SRRM2 (*19*). Our study provides compelling evidence showcasing the detrimental effect of CTCF/cohesin disruption on the association between chromatin and nuclear speckles, resulting in impaired induction of genes located in close proximity to speckles, while having minimal impact on genes distantly located from speckles (**Fig. 4**), supporting a pivotal role of nuclear speckle association in facilitating gene induction. Furthermore, our analyses suggest that, while CTCF’s involvement in enhancer-promoter interactions affects specific genes, its broader impact on gene inducibility may stem from its role in mediating ground-state speckle association (see **Fig. S3E**). More detailed separation of function studies is needed to distinguish between the loop and speckle relevant functions of CTCF and cohesin complex.

We identified a potential Speckle Targeting Motif (STM) within the Rad21 protein (**Fig. 3K**). This 15-amino acid sequence shares a proline periodicity similar to two other factors implicated in nuclear speckle targeting, p53 (*7*) and HIF2A (*20*). Our recent study (*20*) also identifies as much as 1755 proteins harboring putative STM, all of which could contribute to the association between chromatin and nuclear speckle. While the mechanisms behind STM-mediated speckle association are still unclear and warrant additional investigation, our findings offer a molecular basis for future in-depth exploration into the workings of STMs and their interactions with other molecular functions within gene regulatory pathways.

Our observations, in conjunction with previous studies, underscores the significance of nuclear speckles as vital hubs for genes that necessitate high and/or rapid induction. Disruption of this hub could lead to aberrant gene expression patterns and potentially contribute to disease outcomes. Our findings highlight the role of RAD21 variation in causing speckle dissociation and impairing gene induction in CdLS patient-derived cells (**Fig. 5**). Moreover, multiple reports have implicated mutations within nuclear speckle components, including SON (*53*) and SRRM2 (*54*), in neurodevelopmental diseases. It was also discovered that CTCF, SON, and SRRM2 were all among the 285 genes most strongly associated with developmental disorders (*55*). Further exploration of nuclear speckles could thus shed light on the molecular mechanisms underlying gene dysregulation in diseases and uncover potential therapeutic targets.

## Acknowledgments

We thank Dr. Jan-Michael Peters for kindly providing us with the CTCF-AID and RAD21-AID HeLa cell lines.

## Author contributions

Conceptualization: RY, KAA, SLB

Methodology: RY, SR, APS, SCN, KAA

Investigation: RY, SR, APS

Visualization: RY

Funding acquisition: EFJ, BBL, IDK, SLB

Project administration: RY, SLB

Supervision: EFJ, BL, IDK, SLB

Writing – original draft: RY, KAA, SLB

Writing – review & editing: RY, KAA, SLB, EFJ, BBL

## Data and materials availability

Plasmids generated from this study are available from the corresponding author upon request. The genomic datasets generated during this study will be deposited onto GEO. Reviewer links will be provided upon request.

The imaging data supporting this study have not been deposited in a public repository due to file sizes but are available from the corresponding author upon request.

The software, instructions, and code generated by this study used for DNA-FISH analysis and analysis of speckle characteristics are available at https://github.com/Chalietia/CellProfiler

The instructions and code used for analysis of SON TSA-seq data is available at: https://github.com/katealexander/TSAseq-Alexander2020/tree/master/genomicBins_DiffBind

## List of Supplementary Materials

Materials and Methods

Fig.1 to 5, S1 to S4

## Materials and Methods

### 1. Experimental methods

#### a. Cell culture and gene knockdown

HeLa CTCF-AID and HeLa RAD21-AID cell lines constructed in previously work (*24*) were grown in DMEM high glucose 4.5 gm/L - no sodium pyruvate (Corning, 10-017-CM) supplemented with 10% fetal calf serum and 1% Pen/Strep. CTCF knockdown was induced by 24 hours of 500uM Indole-3-acetic acid sodium (Sigma-Aldrich I5148), and control cells treated with 0.1% DMSO unless otherwise noted. siRNA targeting either *WAPL* (siRNA 148512, AM16708, ThermoFisher) or negative control (AllStars Negative Control siRNA, QIAGEN) was transfection with DharmaFECT (FisherScientific, T-2001-02) according to manufacturer protocol and incubated for 48 hours prior to harvesting.

Passage 25-35 IMR90 primary lung fibroblast cell line acquired from ATCC (Cat#CL186) were grown in DMEM (ThermoFisher) supplemented with 10% fetal calf serum and 1% Pen/Strep. siRNA targeting either *CTCF* (Hs_CTCF_6 FlexiTube siRNA, QIAGEN) or negative control (AllStars Negative Control siRNA, QIAGEN) was transfection with DharmaFECT (FisherScientific, T-2001-02) according to manufacturer protocol and incubated for 48 hours prior to harvesting.

HCT116 HaloTAG-CTCF cells were cultured in McCoy media (Thermo Fisher Scientific, 16600082) with 10% FBS (Peak Serum, PS-FB2) and 1%Pen/Strep (Life Technologies, 15070063). CTCF knockdown was induced for 32 hours with HaloPROTAC3 (330nM, AOBIOUS AOB36136) or DMSO control (0.1% DMSO).

Lymphoblastoid cell lines were cultured in RPMI 1640 with 2mM L-glutamine (Invitrogen, 11875085) supplemented with 20% fetal calf serum and 1% Pen/Strep.

#### b. Western blotting and antibodies

Samples were lysed with RIPA buffer (150nM NaCl, 1% IGEPAL, 0.5% sodium deoxycholate, 0.1% SDS and 50mM Tris pH7.4). Protein concentration was determined by bicinchoninic acid assay (BCA) protein assay (#23227, Life Technologies). Samples were separated using precast 4-12% Bis-Tris polyacrylamide gels (ThermoFisher Scientific, NP0321BOX) in the presence of SDS. Proteins were transferred to nitrocellulose membrane, blocked with 5% milk TBST and probed with the following primary antibodies overnight at 4°C: 1. Anti-CTCF antibody (1:5000, abcam, ab188408); 2. Anti-RAD21 antibody (1:1000, Cell Signaling, 4321s); 3. Anti-GAPDH antibody (1:1000, Cell Signaling, 5174); 4. Anti-H3 antibody (1:2500, abcam, ab1791). Following TBST wash steps, membranes were incubated in secondary goat anti rabbit antibody (1:10000, Bio-Rad, 1706515) for one hour, washed with TBST, and detected using using Thermo Scientific SuperSignal West Pico PLUS Chemiluminescent Substrate (ThermoFisher Scientific, 34577). Signal quantification was performed using ImageJ software.

#### c. Immunofluorescence and antibodies

Cells were washed twice with PBS and fixed with Paraformaldehyde solution 4% in PBS (Santa Cruz Biotechnology, CAS 30525-89-4) for 10 minutes at room temperature, washed again with PBS (Corning, 21-031-CV) then permeated with 0.1% PBS Triton X-100 for 15 minutes. Cells were incubated overnight in primary antibodies targeting GFP to stain GFP labelled CTCF or RAD21 (Roche, 11814460001) or SON (abcam, ab121759) diluted 1:200 in PBS at 4C. Secondary antibody conjugated with Goat anti-Rabbit IgG Alexa Fluor™ 488 (ThermoFisher Scientific, A-11008) and Goat anti-Mouse IgG Alexa Fluor™ 647 (ThermoFisher Scientific, A-21235) were diluted 1:200 in PBS and used respectively. After 1 hour incubation at room temperature, samples were washed and incubated in 1:10000 DAPI (Invitrogen, D1306, 5 mg/mL) for 5min. All imaging in this study was performed with Nikon Eclipse Ti2 widefield microscope. For IF only experiments, cells were imaged at focal plane with a 60x oil-immersion objective, and a deep depletion CCD camera cooled to between -70 and −80C.

#### d. ImmunoDNA-FISH

Cells were treated with same protocol as immunofluorescence and same antibodies as IF. After secondary antibody incubation, slides were fixed with 4% PFA in PBS for 10 minutes at room temperature, washed with PBS and permeabilized with 0.5% Triton in PBS for 15 minutes. Subsequently, DNA-FISH was performed as described (*56*). Oligopaint DNA-FISH probes to target genes were designed across a 50kb region centered on the transcription start site as described (*57*). Cells were imaged in a series of 28 optical sections spaced 0.286 microns apart with a 60x oil-immersion objective.

#### e. SON Cut & Run

SON Cut & Run were performed as previously described with minor modification (*21*). For each condition, nuclei from ∼600k cells was isolated using nuclear isolation buffer [10mM HEPES-KOH (pH 7.9), 10mM KCl, 0.1% NP40, 0.5mM spermidine, 1x Halt protease inhibitor cocktail] and bound to concanavalin A lectin beads which had been washed in binding buffer [BioMag Plus; binding buffer: 20mM HEPES-KOH (pH 7.9), 10mM KCl, 1mM CaCl2, 1mMnCl2], as described. After bead binding, samples were split into two tubes, one for each antibody, where binding buffer was replaced with 1:100 antibody diluted in blocking buffer [20mM HEPES-KOH (pH 7.5), 150mM NaCl, 0.1% BSA, 0.5mM spermidine, 1x Halt protease inhibitor cocktail, 2mM EDTA](IgG antibody: Millipore 06-371); SON antibody: abcam ab121759) and incubated overnight at 4°C. Samples were washed in washing buffer [20mM HEPES-KOH (pH 7.5), 150mM NaCl, 0.1% BSA, 0.5mM spermidine, 1x Halt protease inhibitor cocktail], and treated with pA-MNase and CaCl2 for 30 minutes on an ice cold pre-chilled metal block on ice. The reaction was stopped by addition of STOP buffer [200mM NaCl, 20mM EDTA, 4mM EGTA, 50µg/mL RNase A, 40µg/mL glycogen] while gently vortexing, and DNA fragments were released by incubation at 37°C for 10 minutes. The DNA fragments were collected from bead supernatants, treated with Proteinase K for 10 minutes at 70°C, then subject to PCI-choroform purification. Libraries from DNA fragments were made using the NEBNext Ultra II DNA Library Prep Kit for Illumina. Library sizes were determined on a Bioanalyzer, and concentration determined using NEBNext Lib Quant Kit (E7630, NEB). Libraries were sequenced with an Illumina NextSeq550 with 42bp per read (total of 84bp) using the NextSeq 500/550 High Output 75-cycle v2.5 kit.

#### f. SON Cut & Tag

SON Cut&Tag was performed according to the previously published protocol (*58*) with minor modification. For each experiment, 1 million cells were harvested, washed and lysed on ice for 10 minutes. Nuclei were resuspended in PBS and fixed with 0.1% formaldehyde for 2 minutes at room temperature and quenched with 75mM glycine. 250,000 nuclei per condition (as determined by a hematocytometer) were used per condition. Samples were incubated with 1:50 of anti-SON antibody (abcam, ab121759) at 4°C overnight. Beads were then resuspended in secondary antibody in wash buffer (1:100 guinea pig anti-rabbit ABIN101961) and nutated at room temperature for 1 hour. Liquid was removed and beads were washed three times with wash buffer. pA-Tn5 (purified in-house according to published protocol (Priore et al. 2021)) was mixed 1:50 in Dig-300 buffer (without digitonin, 20 mM HEPES-NaOH pH 7.5, 300 mM NaCl, 0.5mM spermidine, Roche Complete Protease Inhibitor EDTA-free) and beads were resuspended 100 uL and nutated at room temperature for 1 hour. Beads were washed three times in Dig595 300 buffer. Beads were resuspended in 300 uL Tagmentation Buffer (Dig-300 buffer with 10 mM MgCl2) and incubated at 37 °C for one hour. 16 mM EDTA, 0.1% SDS, and 50 µg of Proteinase (Invitrogen, 25530015) was added per sample and incubated at 55 °C for one hour. DNA was purified by PCI-chloroform extraction. The purified DNA was then amplified by PCR using NEBNext HiFi 2x PCR Master Mix (NEB M0541S), and the resulting library was purified using 1.3X SPRI purification (Omega Bio-Tek, M1378-01). Libraries were sequenced using a NextSeq1000 (Illumina) with 50bp paired end reads at a sequencing depth of approximately 10-20 million reads per sample.

#### g. Plasmid construction and transfection

Plasmid mutagenesis was performed on pKS070 - pCAGGS-3XFLAG-(human) CTCF-eGFP (Addgene Plasmid #156448) for CTCF and mEmerald-RAD21-N-18 (Addgene Plasmid #54248) for RAD21. Mutagenesis was carried out with QuickChange II Site-Directed Mutagenesis Kit (Agilent, 200523) with primers listed below.

Plasmid transfection was carried out with XtremeGENE HP (Millipore Sigma, 6366244001) according to manufacturer protocol at a reagent/DNA ratio of 2:1 in serum free media. Cells were incubated in transfection mixture for 4 hours then changed to regular media, followed by designated treatments the next day.

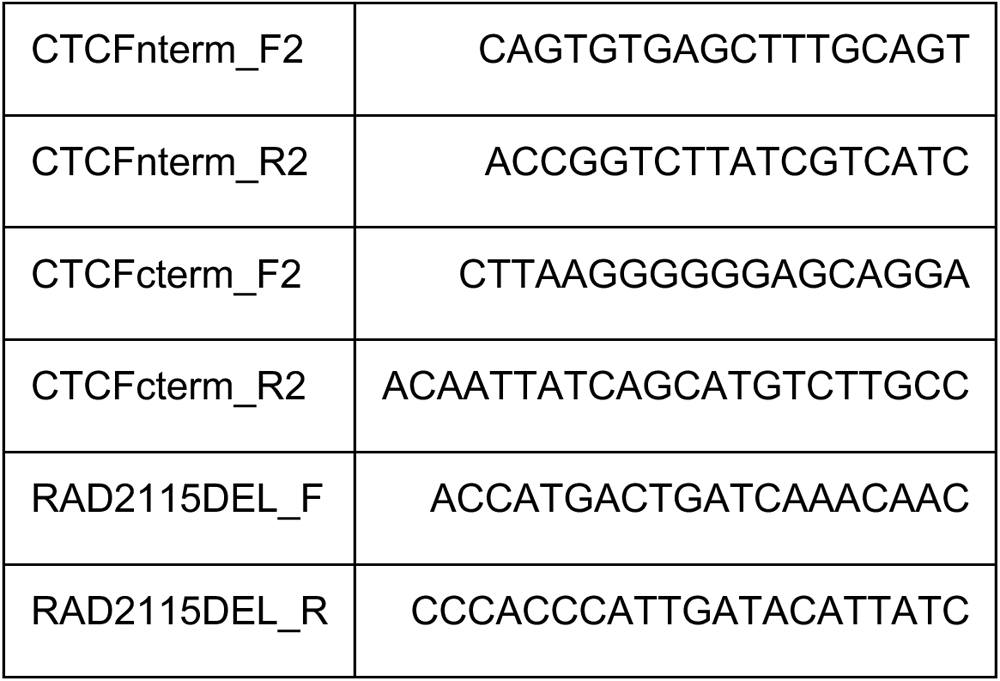

#### h. Polyadenylated RNA-seq

Cells were lysed in TRIzol (15596018, Thermo Fisher Scientific) and snap frozen. RNA was then isolated using chloroform extraction, followed by QIAGEN RNeasy Mini Kit isolation (#74106), including DNA digestion with DNase. Poly(A)+ RNA was further isolated using double selection with poly-dT beads (E7490, NEB). RNA-seq libraries were prepared using the NEBNext Ultra II Directional Library Prep Kit for Illumina (E7760, NEB). The size of the libraries was determined using a Bioanalyzer, and their concentration was determined using the NEBNext Lib Quant Kit (E7630, NEB). The libraries were sequenced on an Illumina NextSeq 550, utilizing paired-end sequencing with 42 bases per read, at a sequencing depth of approximately 15 million reads per sample.

### 2. Data processing and analysis

#### a. ChIP-seq correlational and permutation analysis

Spearman correlation in 1A was calculated by deepTools *multiBigwigSummary* with bin size (-bs argument) of 50000. All previously published genomic resources used in this study are listed in Table S3 (3. Genomics data reference).

Permutation analysis in 1C was performed as follow: 1. List of predicted binding target genes of designated factors were retrieved from Transcription Factor Gene Database (https://tfbsdb.systemsbiology.net/). Motif list used are listed as follow, CTCF: CTCF_C2H2_full_monomeric; YY1: YY1_C2H2_full_monomeric; p53: Tp53_p53l_DBD_dimeric; AP1: V_AP1_01_M00517. 2. Genomic coordinates of target gene list spanning +50kb and -50kb of transcription start site (TSS) were used as query due to the broad distribution of SON signal across the genome. 3. Permutation tests with RegioneR (https://bioconductor.org/packages/release/bioc/html/regioneR.html) were performed using genomic regions spanning 50 kilobases upstream and downstream of all hg19 RefSeq gene transcription start sites as the background (universe), *with randomize. function= resampleRegions, evaluate.function=meanInRegions*, and randomize time *n=1000*.

#### b. Immunofluorescence (IF) data processing and analysis

All IF images were taken at the focal plane with the autofocus function of Nikon NIS software on a Nikon Eclipse Ti2 widefield microscope with a 60x oil-immersion objective lens. Subsequent image processing was performed with CellProfiler (https://cellprofiler.org/) to identify nuclear speckle and measure various size and shape parameters. Pipeline and notes are available on Github: https://github.com/Chalietia/CellProfiler/blob/main/SpeckleSPEC1.1.cppipe

#### c. ImmunoDNA-FISH data processing and analysis

For All ImmunoDNA-FISH, cells were imaged in a series of 28 optical sections. Resulting image stacks were first maximum projected with *MakeProjection* function of CellProfiler. All DNA-FISH foci within DAPI stained nucleus were identified, nucleus with more than 4 identified FISH foci were removed. After identifying nuclear speckle, the distance between center of FISH foci and the edge of its nearest nuclear speckle was measured in pixel. In subsequent post-processing, pixel distance* 0.11 um/pixel was calculated as the final physical distance. Pipeline, QC and sample data are available on https://github.com/Chalietia/CellProfiler. Speckle distance distribution p values were calculated using Mann-Whitney-Wilcoxon tests.

#### d. SON Cut&Run/SON Cut&Tag data processing

Sequencing reads were aligned to human reference genome assembly GRC37/hg19 with the following parameter using Bowtie2(*59*): --local --end-to-end --very-sensitive --no-mixed --no-discordant --phred33 -I 10 -X 1000-x hg19, subsequently filtered with samtools view -q 5 and deduplicated with Picard (https://broadinstitute.github.io/picard/). According to ours and previous observation (*7*, *9–11*), SON signal across genome is broad and not as highly enriched as regular transcription factors. Therefore, to minimize the impact of noise, sharp peaks were called for all bam files by macs3 callpeak -f BAM -g 2.7e9 -q 0.1. The noise peaks have an average size of 200bp and cover less than 0.2% of the effective genome. Combined noise peaks files were then merged with ENCODE hg19 blacklist (https://github.com/Boyle-Lab/Blacklist/blob/master/lists/hg19-blacklist.v2.bed.gz) and excluded from all subsequent analysis. For all results except CTCF binding region visualization in Fig. 2G-H, Bigwig and BedGraph files were generated with deepTools: bamCompare -b1 $SON -b2 $IGG -bs 5000 --smoothLength 15000 --effectiveGenomeSize 2864785220 --exactScaling --normalizeUsing RPKM --scaleFactorsMethod None -bl blacklistPEAK.merge.bed. For Fig. 2G-H, given the narrower nature of CTCF binding peaks, Bigwig files were generated with -b $SON -bs 50 -- effectiveGenomeSize 2864785220 --exactScaling --normalizeUsing RPKM.

#### e. SON Cut&Run QC

For correlation between SON TSA-Seq and SON Cut&Run, average IgG normalized SON signal from both experiments was calculated and compared across CDS of all RefSeq genes. For comparison between MERFISH and SON Cut&Run, MERFISH data containing location and distance between probes and nearest nuclear speckle was retrieved from https://zenodo.org/record/3928890. Average IgG normalized SON signal at corresponding genomic loci was calculated by summing up signal in corresponding bedgraph files and compared. The top 5% most distant loci in MERFISH data were considered as outlier and excluded from this analysis.

For comparison between ImmunoDNA-FISH and SON Cut&Run, average IgG-normalized SON signal from TSS -25kb to TSS +25kb of corresponding genes was calculated and compared. This region is where the Oligopaint DNA-FISH probes were located.

#### f. Speckle associated domains (SPADs) calling

SPAD calling was performed by SICER (*60*) with the following parameters: sicer -t $SON -c $IGG -s hg19 -fdr 0.1 -w 2000 -g 20000. To identify consensus regions, SPADs identified from two technical replicates for each condition were intersected as final output.

#### g. SON Cut&Run differential analysis

SON Cut&Run differential analysis was based on previously published method (*7*) (step-by-step instruction available at https://github.com/katealexander/TSAseq-Alexander2020/tree/master/genomicBins_DiffBind) with minor modification. Briefly, from bam files with PCR duplicates removed, SON signal was quantified over sliding windows of 50kb (slid by 1/10th of the window size) using DiffBind (https://bioconductor.org/packages/release/bioc/html/DiffBind.html) with input subtraction. Blacklist files generated in (d) were set as DiffBind blacklist with dba.blacklist(). The normalization method was set to DBA_NORM_RLE. Regions with significant increase or decrease of SON signal were separated into different files, and those regions with FDR<0.1 was considered as differential and merged using BEDTools. Genes of which TSS located within differential domains were extracted using a Python script, as in https://github.com/katealexander/TSAseq-Alexander2020/blob/master/genomicBins_DiffBind/getGenesWithin.py.

#### h. Gene ontology (GO) analysis

Genes exhibiting the highest 10% SON Cut&Run signal across coding sequences (CDS) in both IMR90 and HeLa cells were selected as input for the GO Biological Process analysis with default setting on STRING database (http://string-db.org). Resulting GO categories with strength>0.2, fdr<0.01, and a background 100 to 3000 genes were subjected to REVIGO (http://revigo.irb.hr/) analysis to identify non-overlapping categories. The GO terms with largest background within each cluster were selected for visualization.

#### i. RNA-seq data processing

RNA-seq data was aligned to the reference human genome assembly, GRC37/hg19, using the splice-aware STAR alignment (https://github.com/alexdobin/STAR/releases) with the following settings: --outFilterType BySJout --outFilterMultimapNmax 20 –alignSJoverhangMin 8 --alignSJDBoverhangMin 1 --outFilterMismatchNmax 999 -- outFilterMismatchNoverReadLmax 0.04 --alignIntronMin 20 --alignIntronMax 1000000. Reads for each gene with RefSeq annotations were counted using htseq-counts (*61*). Normalized counts and differential analysis were performed with Deseq2 (*41*). Statistical differences between log2FC in different SON signal bins were determined using Mann-Whitney-Wilcoxon tests.

#### j. RNA-seq interaction analysis

Interaction analysis was performed based on Deseq2 instruction (https://www.bioconductor.org/packages/devel/bioc/vignettes/DESeq2/inst/doc/DESeq2.html <u>#interactions</u>), with the following design: design= ∼genotype + condition + genotype: condition. To specifically look at how condition effect (e.g., heat shock) differ across genotype (e.g., CTCF KD), the interaction term was extracted with results (dds, name=“genotypeII.conditionB”). All genes with a p value <0.05 were considered to have significantly differential conditional effect.

#### k. Categorization of loop status

For calling CTCF sites involved in significant loop or otherwise, CTCF ChIP-seq and HiChIP data in HeLa cell were obtained from GSE108869. CTCF binding peak was called by MACS2 with default parameter with q<0.01 as original publication. For HiChIP, combined loop file from 2 replicates of CTCF HiChIP experiments were subjected to HICDC+ for loop calling with a binsize=10kb and all interactions with q value<0.01 were considered significant. Identified CTCF peaks located within any significant bin were considered to be looping CTCF sites, and those do not overlap with significant bin were considered non-looping CTCF sites. Genes of which TSS overlaps with CTCF sites were identified and separated based on looping status. Subsequently, we chose the highest 20% of CTCF-bound genes proximal or distal from nuclear speckles and analyzed gene induction changes under heat shock as in (j).

### 3. Genomics data reference

All previously published data used in this study is listed below:

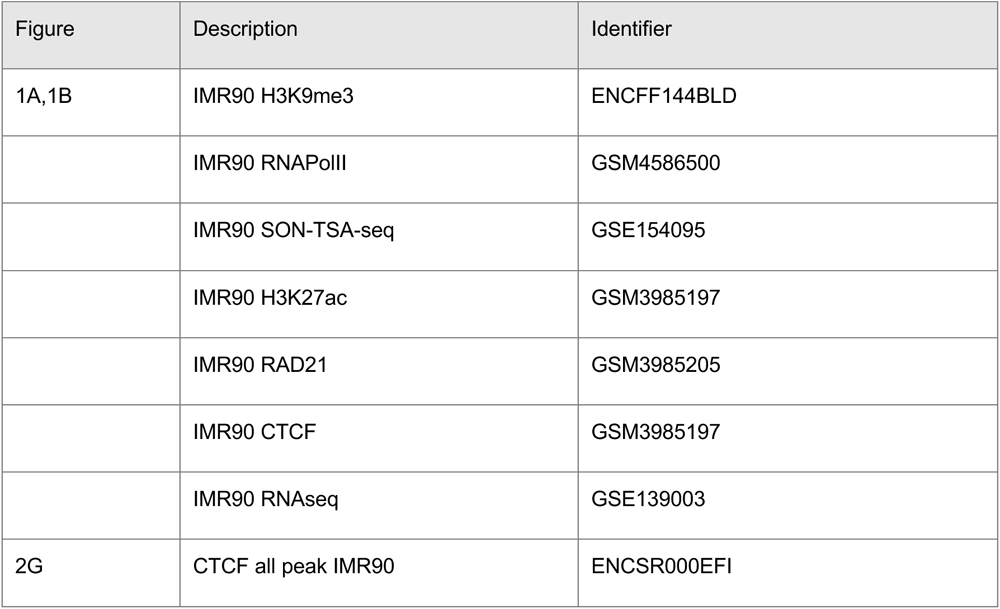
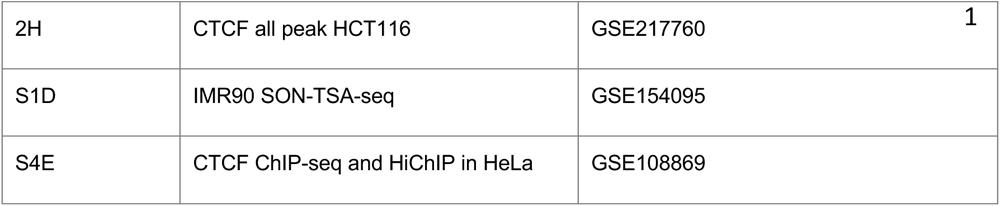

---

**Figure S1.**
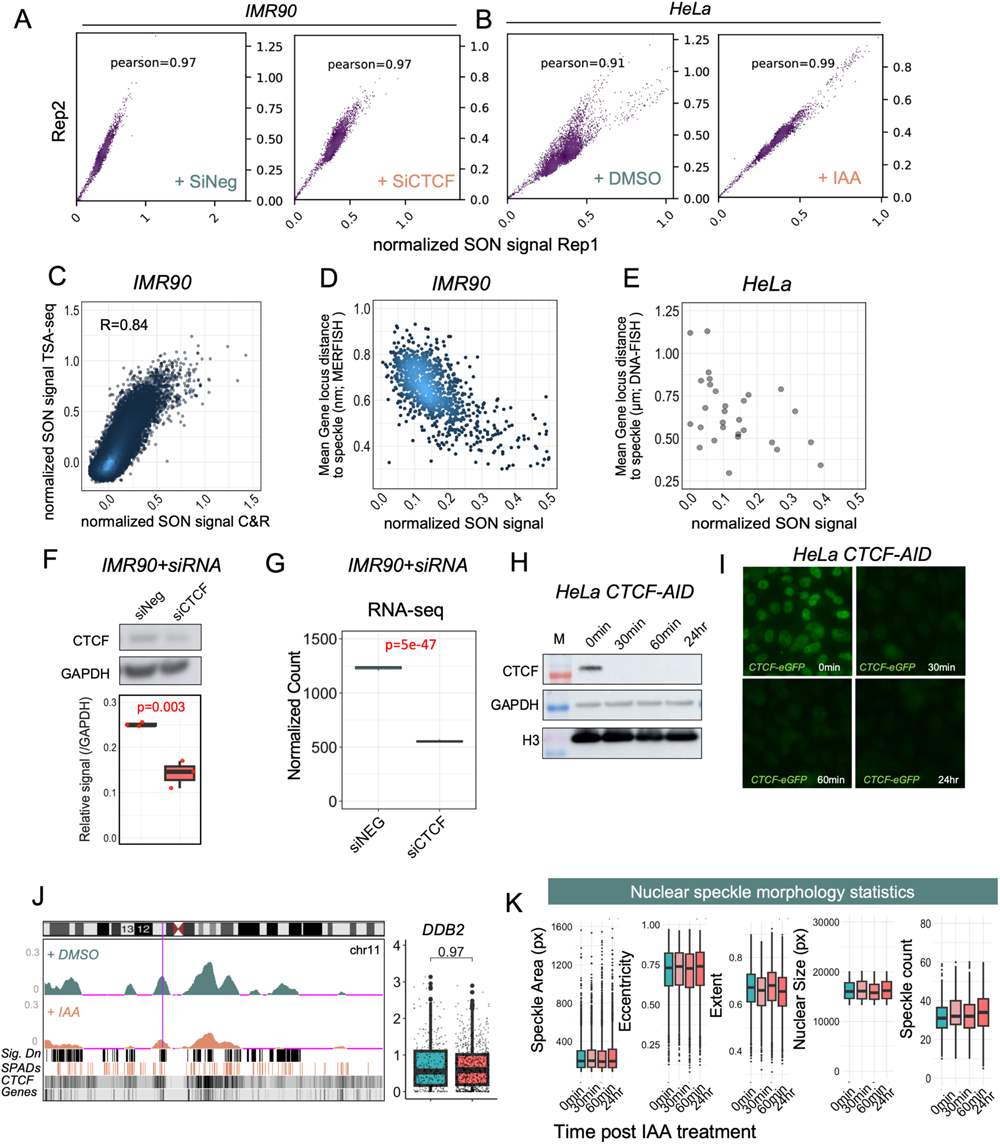
Quality control of CTCF depletion experiments. A. Correlation between technical duplicates of designated SON Cut&Run performed in IMR90 cells. B. Correlation between technical duplicates of designated SON Cut&Run performed in HeLa cells. C. Correlation between SON TSA-seq and SON Cut&Run signal over all CDS across genome in IMR90 cells. D. Correlation between nuclear speckle proximity measured by SON Cut&Run and MER-FISH in IMR90 cells. E. Correlation between all nuclear speckle proximity data measured by DNA-FISH and SON Cut&Run under same condition obtained in this study. F. Western blot against CTCF in IMR90 cell line with siRNA against negative control or CTCF. (Top) Representative blot, (bottom) quantification. G. RNA-seq normalized counts of designated samples. H. Western blot against CTCF in CTCF-AID HeLa cells, at different timepoints after auxin treatment. M represents marker. I. Immunofluorescence against CTCF-eGFP (green) at different timepoints after auxin treatment. J. (Left) Whole chromosome 11 view of SON Cut&Run data under designated treatment conditions. (Right) Quantification of DDB2 gene loci distance to nearest nuclear speckle upon DMSO (green) or auxin (red) treatment in CTCF-AID HeLa cell lines. K. Morphological traits of nuclear speckles measured from Immunofluorescence against speckle marker SON, at designated timepoints after auxin treatment.

**Figure S2.**
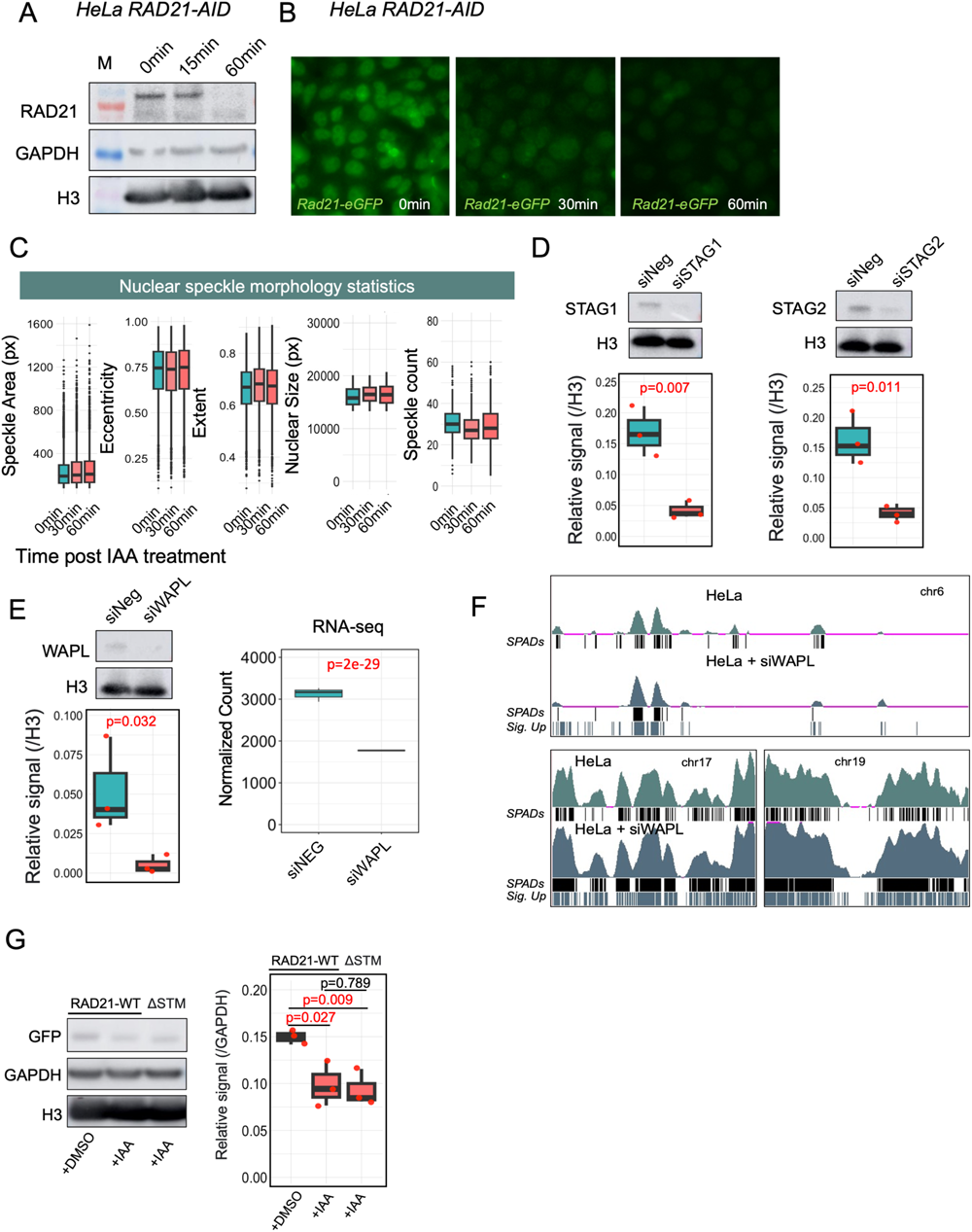
Quality control of RAD21 depletion experiments. A. Western blot against RAD21 in RAD21-AID HeLa cells, at different timepoints after auxin treatment. M represents marker. B. Immunofluorescence against RAD21-eGFP (green) at different timepoints after auxin treatment. C. Morphological traits of nuclear speckles measured from Immunofluorescence against speckle marker SON, at different timepoints after auxin treatment in RAD21-AID HeLa cells. D. (Left) Western blot against STAG1 in HeLa cell line with siRNA against negative control or STAG1. (Top) Representative blot, (bottom) quantification. (Right) Western blot against STAG2 in HeLa cell line with siRNA against negative control or STAG2. (Top) Representative blot, (bottom) quantification. E. (Left) Western blot against WAPL in HeLa cell line with siRNA against negative control or WAPL. (Top) Representative blot, (bottom) quantification. (Right) RNA-seq normalized counts of designated samples. F. Whole chromosome 6,17,19 view of SON Cut&Run data in HeLa cells with siRNA against WAPL or negative control for 48hr. G. Western blot against RAD21-GFP in HeLa cell line with designated transfected construct. (Left) Representative blot, (right) quantification.

**Figure S3.**
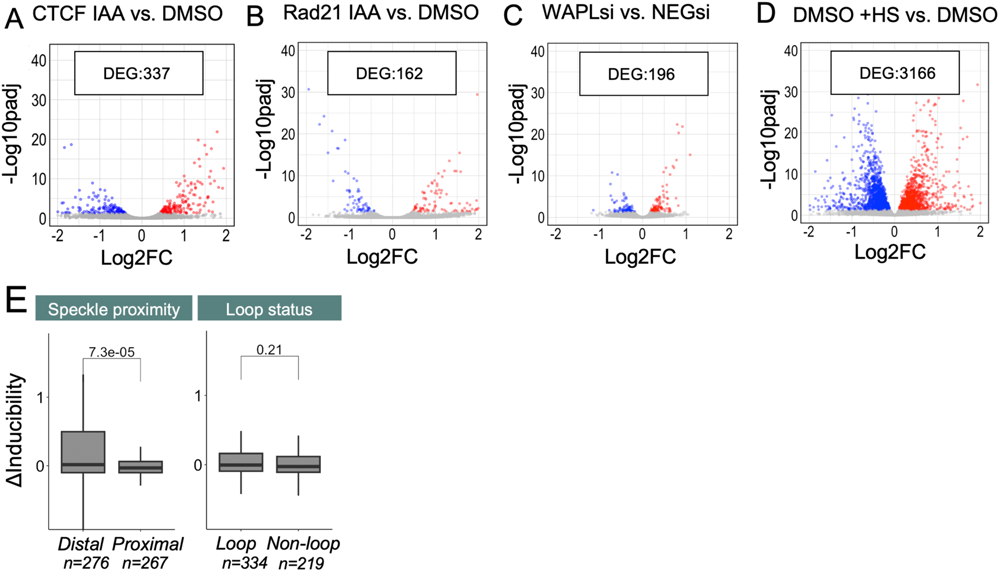
Disruption of nuclear speckle association hinders the induction of nuclear speckle proximal genes. A. Differential expression analysis of HeLa cells with or without 24hr auxin treatment to degrade CTCF. Blue dots represent genes significantly down-regulated (Log2 fold change <0, p<0.01), red dots represent genes significantly up-regulated (Log2 fold change >0, p<0.01). DEG: deferentially expressed genes. B. Differential expression analysis of HeLa cells with or without 1hr auxin treatment to degrade RAD21. C. Differential expression analysis of HeLa cells with or without 48hr siRNA mediated degradation of WAPL. D. Differential expression analysis of HeLa cells subjected to a 1-hour heat shock treatment, with or without a 24-hour auxin treatment to degrade CTCF. E. Comparison of conditional effect (ΔInducibility) between auxin induced CTCF KD and DMSO treated HeLa cells under heat shock. Genes with significant differential conditional effect were separated based on speckle proximity (Left) or whether they locate proximal (within 10kb) to loop anchors.

**Figure S4.**
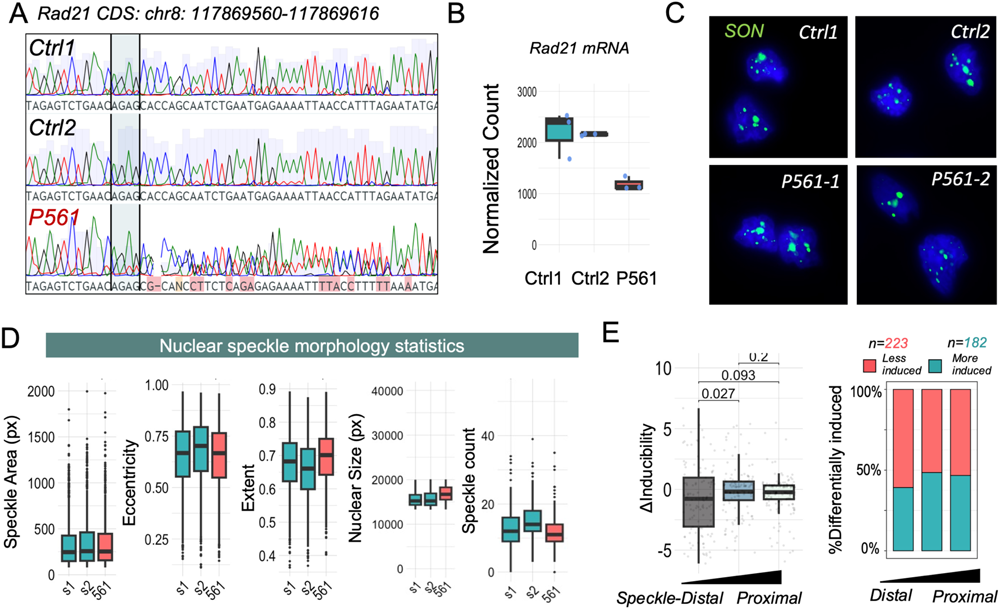
Quality control and transcriptomic profiling of donor derived LCLs. A. Sanger sequencing results of designated derived LCL cells, within RAD21 gene coding region at chr8: 117869560-117869616. B. RNA-seq normalized counts of RAD21 genes in designated healthy individual or patient derived LCLs. C. Representative images with DAPI stained nuclei (blue), immunofluorescence of nuclear speckle component SON (green), in LCL cells from healthy donors (Ctrl1, Ctrl2) and patient 561 (P561-1, P561-2). D. Morphological traits of nuclear speckles measured from immunofluorescence against speckle marker SON in designated healthy individual or patient derived LCLs. E. Differential conditional effect under heat shock between two control LCLs from healthy individual.

